# An ABC method for whole-genome sequence data: inferring paleolithic and neolithic human expansions

**DOI:** 10.1101/419002

**Authors:** Flora Jay, Simon Boitard, Frédéric Austerlitz

**Author notes:** Corresponding author: Flora Jay.

## Abstract

Species generally undergo a complex demographic history, consisting, in particular, of multiple changes in population size. Genome-wide sequencing data are potentially highly informative for reconstructing this demographic history. A crucial point is to extract the relevant information from these very large datasets. Here we designed an approach for inferring past demographic events from a moderate number of fully sequenced genomes. Our new approach uses Approximate Bayesian Computation (ABC), a simulation-based statistical framework that allows (i) identifying the best demographic scenario among several competing scenarios, and (ii) estimating the best-fitting parameters under the chosen scenario. ABC relies on the computation of summary statistics. Using a cross-validation approach, we showed that statistics such as the lengths of haplotypes shared between individuals, or the decay of linkage disequilibrium with distance, can be combined with classical statistics (eg heterozygosity, Tajima’s D) to accurately infer complex demographic scenarios including bottlenecks and expansion periods. We also demonstrated the importance of simultaneously estimating the genotyping error rate. Applying our method on genome-wide human-sequence databases, we finally showed that a model consisting in a bottleneck followed by a Paleolithic and a Neolithic expansion was the most relevant for Eurasian populations.

## Introduction

The inference of demographic history from genetic polymorphism data is a long-standing subject in population genetics (Veeramah and Hammer 2014; Schraiber and Akey 2015). Since high-throughput sequencing data are becoming increasingly available, it becomes imperative to develop novel methods aiming at handling these large amounts of data of varying quality.

There are currently several kinds of methods available. The most common ones are based either on the repartition of polymorphic sites along the genome or on the analysis of the site frequency spectrum (SFS). The first class includes (i) Hidden Markov model (HMM) methods based on the sequential Markov coalescent (SMC and SMC’) (McVean and Cardin 2005; Marjoram and Wall 2006) such as PSMC (Li and Durbin 2011), MSMC (Schiffels and Durbin 2014), diCal (Sheehan et al. 2013), and SMC++ (Terhorst et al. 2017), and (ii) methods using the lengths of regions that are identical-by-state (IBS) or identical-by-descent (IBD) within pairs of haplotypes (Browning and Browning 2015; Palamara et al. 2012; Harris and Nielsen 2013; MacLeod et al. 2013). These methods capture the recombination process and at least partial knowledge on the hidden genealogies, and therefore allow extracting substantial information even from a very low number of individuals (as low as 1 for PSMC). A strong limitation of HMM-based methods is that increasing this number highly increases the complexity of analytical results and the amount of computational power required, making them unusable for larger samples (usually above five individuals). This impedes the inference of recent demographic events (Schiffels and Durbin 2014; Boitard et al. 2016). Note, however, that the more recent SMC++ scales to a large number of individuals, and thus more recent time scales, by combining information from TMRCAs of all haplotype pairs and the SFS (Terhorst et al. 2017). The IBD-based methods on the other hand retrieve efficiently recent events but cannot infer ancient history (Browning and Browning 2015).

The second class of methods is based on the analytical analysis of the site frequency spectrum and assumes independent segregating sites (e.g. Gutenkunst et al. 2009; Bhaskar et al. 2015; Liu and Fu 2015). These methods, which do not take the recombination process into account, are easily scalable to very large sample size datasets, such as large SNP chips or genome-wide sequences. Because these SNP-based approaches are blind to linkage information, and because it might be hard in practice to distinguish two SFS computed from limited amount of data, they are poorly informative about recent history unless the sample size is quite large (in the order of hundreds, see Bhaskar and Song (2014) for results on identifiability).

Moreover, apart from *dadi* (Gutenkunst et al. 2009), all the methods previously mentioned (SMC, IBD or SFS based, …) do not propose a formal testing between competing demographic models. For this purpose, Approximate Bayesian Computation (ABC) offers a proper framework aiming at investigating several models and selecting the best-fitting one, along with inferring its parameters. ABC is a likelihood-free approach that consists in simulating a large number of pseudo-datasets under several demographic scenarios; the best scenario is then chosen by analyzing which pseudo-datasets are the closest to the observed data (Csilléry et al. 2010; Sunnåker et al. 2013) The parameters of this scenario (e.g. effective population sizes, migration rates, growth rates, split times, …) are then estimated similarly. ABC has been shown to be a valuable method to infer population history, and has been applied widely to many kind of genetic polymorphism data, including microsatellite data, single nucleotide polymorphism (SNP) data, genotype-by-sequencing data, and short autosomal sequences (e.g. Excoffier et al. 2005; Fontaine et al. 2012; Sjödin et al. 2012; Shafer et al. 2014; Palstra et al. 2015).

A few ABC methods or other kinds of simulation-based approaches have been proposed to tackle whole-genome data and investigate complex demographic scenarios. Excoffier et al. (2013) developed a composite-likelihood approach based on simulations, which relies exclusively on the joint site frequency spectrum. Although this method differs from the purely analytical SFS-based methods (Bhaskar et al. 2015; Liu and Fu 2015), it again requires a large sample size to counterbalance the loss of linkage information. Wollstein et al. (2010) developed an ABC approach that uses jointly the SFS and short haplotype diversity computed from DNA chips data. Li and Jakobson (2012) developed another ABC approach based on haplotype information, short-distance linkage disequilibrium and a few traditional population genetics statistics, all computed on 100Kb genomic regions. However, extremely few simulation-based methods are specifically designed for whole-genome sequence data and the practicability of ABC for long recombining sequences (≥2Mb) was investigated only in few studies (Theunert et al. 2012; Boitard et al. 2016). This scarcity is likely due to the computational burden of simulating long regions of the genome and computing complex summary statistics for each pseudo-dataset. Theunert et al. (2012) developed two new statistics: allele frequency-identity by descent (AF-IBD) and allele frequency-identity by state (AF-IBS), which was computed for simulated long-recombining regions and genome-wide SNP data. Boitard et al. (2016) also simulated long-recombining regions and computed both SFS and linkage disequilibrium statistics. These two studies demonstrated that the approach is feasible and that useful information can be extracted from dense data of intermediate sample size. Yet, Theunert et al. (2012) did not apply their method to whole-genome sequencing but to SNP arrays, while Boitard et al. (2016) did not investigate model testing. Moreover, as these studies focused on specific categories of summary statistics, they did not specifically explore which combinations of summary statistics would lead to the best demographic inference, among all the classical statistics proposed in the literature.

Improving existing inference methods is especially relevant for understanding human evolution history. Even if some demographic processes have been reconstructed repeatedly, such as a strong bottleneck in population size for non-African populations resulting from the out-of-Africa migration (Veeramah and Hammer 2014), many other parts of history are still obscure or controversial. For instance, several studies have shown that food-producing human populations increased in size since the Paleolithic, while this was not the case for hunter-gatherer populations (Excoffier and Schneider 1999; Patin et al. 2009; Aime et al. 2013). However, the inferred effective sizes and timings of expansions vary widely across studies and seem sensitive to the inference method and sample size used. While some studies inferred an expansion starting in the upper Paleolithic (Cox et al. 2009; Patin et al. 2009; Batini et al. 2011), other studies using rapidly mutating microsatellite markers (Aimé et al. 2014) or specific mitochondrial haplogroups (Soares et al. 2012) point toward a more recent Neolithic expansion which is consistent with the hypothesis of two subsequent expansions (Tennessen et al. 2012). A recent simulation study (Aimé and Austerlitz 2017) has shown, indeed, that in case of a Paleolithic expansion followed by a Neolithic expansion, slowly mutating markers such as nuclear DNA sequences will detect only the ancient expansion, while rapidly mutating microsatellite markers will only detect the recent one. These studies provide thus an indirect evidence of this two-expansion process, however the question remains whether we can infer this process directly from whole-genome data.

Here, we developed a model-based ABC approach for next-generation sequencing data, and investigated the practicability of using a very large number of summary statistics to maximize the information that can be extracted from a small or intermediate number of sequences. We also investigated the power of the method to infer recent history and especially to detect different phases of growth. We proposed a simple solution to account for sequence data specificities, investigating in particular the best way for handling the genotyping error rate. We then applied our method to several human populations of African, European and Asian ancestries, based on two published datasets (Complete Genomics and 1000Genomes). These allowed us to investigate whether these populations were submitted to a bottleneck during the out-of-Africa period and whether they underwent one or two expansion phases during the Paleolithic and the Neolithic.

## New approaches

Our ABC approach leverages whole-genome sequence data to first select the best demographic model among several proposals, and then co-estimate demographic parameters and a genotyping error rate. It is based on realistic simulations of long recombining chromosomal segments encompassing genotyping errors, and on a set of statistics that aims at summarizing most aspects of NGS data, by capturing dependencies between SNPs at different scales along the chromosome. These summary statistics were divided into five categories (see table 1 and Methods for details): (i) *Classical* statistics: Tajima’s D, 50kb-haplotype heterozygosity, etc.; (ii) the site-frequency-spectrum and total number of SNPs *(SFS)*; (iii) the decay of linkage disequilibrium *(LD)*; (iv) the length of identical-by-state segments across two or more chromosomes *(IBS)*; and (v) the extended length of IBS around a SNP conditioned on its derived allele frequency (allele-frequency identity-by-state, AFIBS; Theunert et al. 2012). These choices were based on a great number of previous studies highlighting that each of these summary statistics contains information about past demographic history (Gutenkunst et al. 2009; Theunert et al. 2012; for a review see Gattepaille et al. 2013; Harris and Nielsen 2013; Patin et al. 2014) We evaluated the performance of each category of statistics alone or combined with other categories, by considering first each category alone in the ABC procedure, then all pairwise combinations, and then all five categories simultaneously.

**Table 1.** Summary statistics used in our study, grouped into five classes.

| Abbrev. | Full | Details | Nb stat. * |
| --- | --- | --- | --- |
| **Classical** | Several statistics widely used in ABC | Expected heterozygosity, 50kb-haplotype heterozygosity, Tajima D [1], ... | 7 |
| **SFS** | Site frequency spectrum | Relative counts (mean, sd)<br>Total number of SNPs | 35 |
| **LD** | Linkage disequilibrium (decay) | r<sup>2</sup> for pairs of SNPs for bins of distance [2-4] | 38 |
| **IBS** | Identity-by-state | Distribution of IBS lengths across 2, 4 or 8 chromosomes [5,6] | 44 |
| **AFIBS** | Allele frequency identity-by-state | IBS lengths conditional on the frequency of a focal SNP (binned per AF) [7] | 32 |

\* Number of statistics for a dataset of 9 diploid individuals [1] Tajima ; [2] Hayes ; [3] Patin ; [4] Boitard et al., 2016 ; [5] Harris and Nielsen, 2013 ; [6] MacLeod et al., 2013 ; [7] Developed by Theunert et al 2012

## Results

### Benchmarking ABC routines

We first benchmarked different ABC routines previously developed for parameter estimation and model selection (Blum et al. 2013; see Materials and Methods; Blum and François 2010; Beaumont et al. 2002; Beaumont 2008). We considered a sample size of 9 individuals per population, because this is the number of sequences we could obtain from high quality public datasets for a reasonably large number of human populations. In a preliminary analysis, we considered two demographic models that encompass either one or two exponential expansions in the last 5,000 generations (fig. 1 top, supplementary table S1), comparing the performances of three methods that estimate the posterior model probabilities for model selection (rejection, multinomial logistic, neural network) and four methods that estimate the posterior distribution of parameters for the selected model (rejection, linear regression, ridge regression, neural network). The misclassification rates for model selection (i.e. the probability of selecting the wrong model among the two proposed) and prediction errors for parameter inference (i.e. a measure of estimation accuracy) were computed through cross-validation with 1000 *pseudo-observed datasets*.

**Fig. 1.**
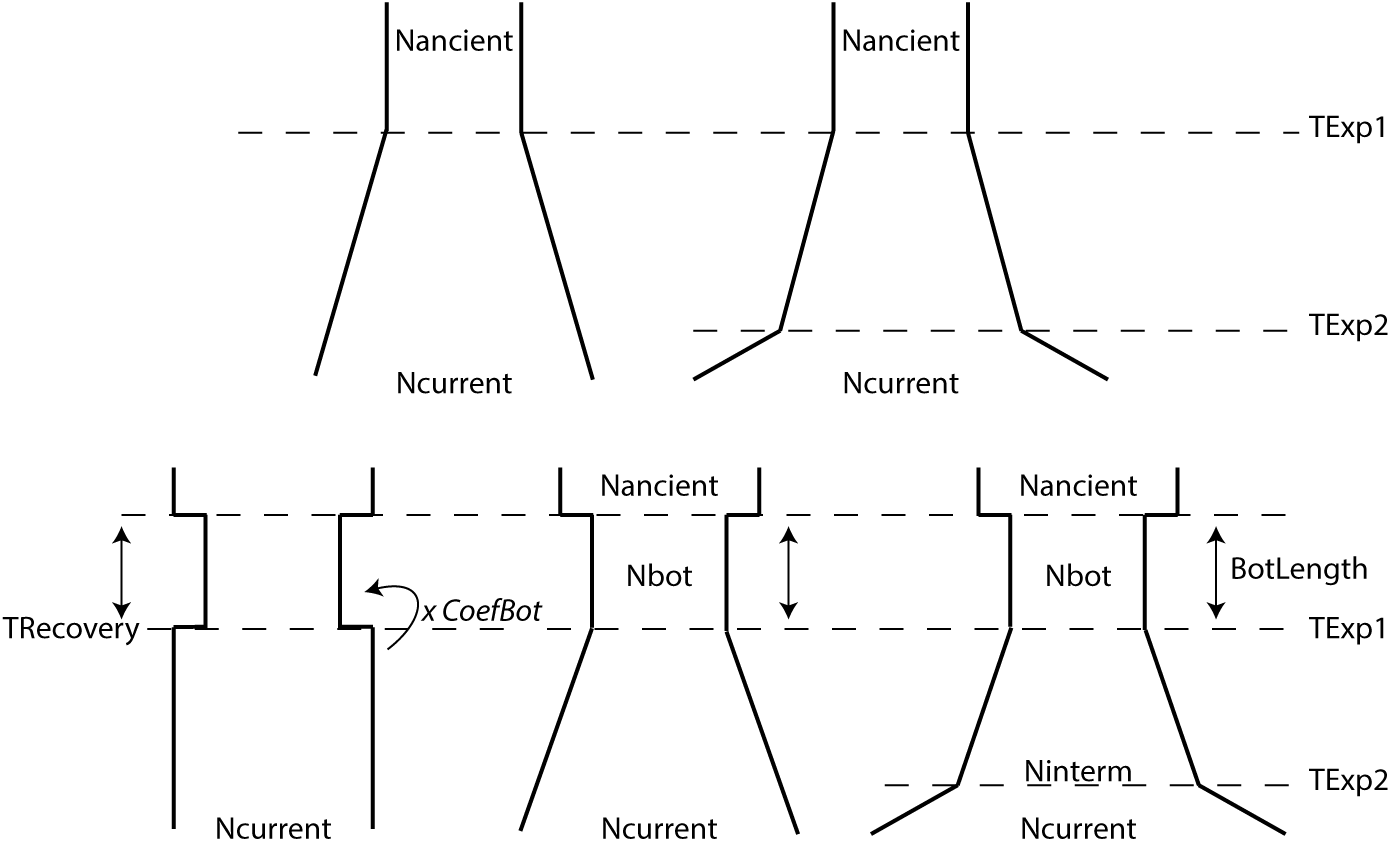
Schemes of the five demographic scenarios considered in this study. The expansions are depicted on a log scale, hence they are all exponential. Details about parameters and prior bounds are described in Supplementary Materials.

The misclassification rate when combining all categories of statistics, was 31% and 10% lower when using the neural network than when using the rejection algorithm and the multinomial regression, respectively (fig. 2A). Similarly, for parameter estimation, for both demographic models and whatever the combination of summary statistics, we found that using neural networks lead to the lowest prediction error: from 30 to 54% lower than when using the basic rejection algorithm (fig. 2B, supplementary table S2). This conclusion is consistent with that of Boitard et al. (2016), who found that neural network regression significantly improved ABC inference of population size histories, when these were modelled by piecewise constant processes.

**Fig. 2.**
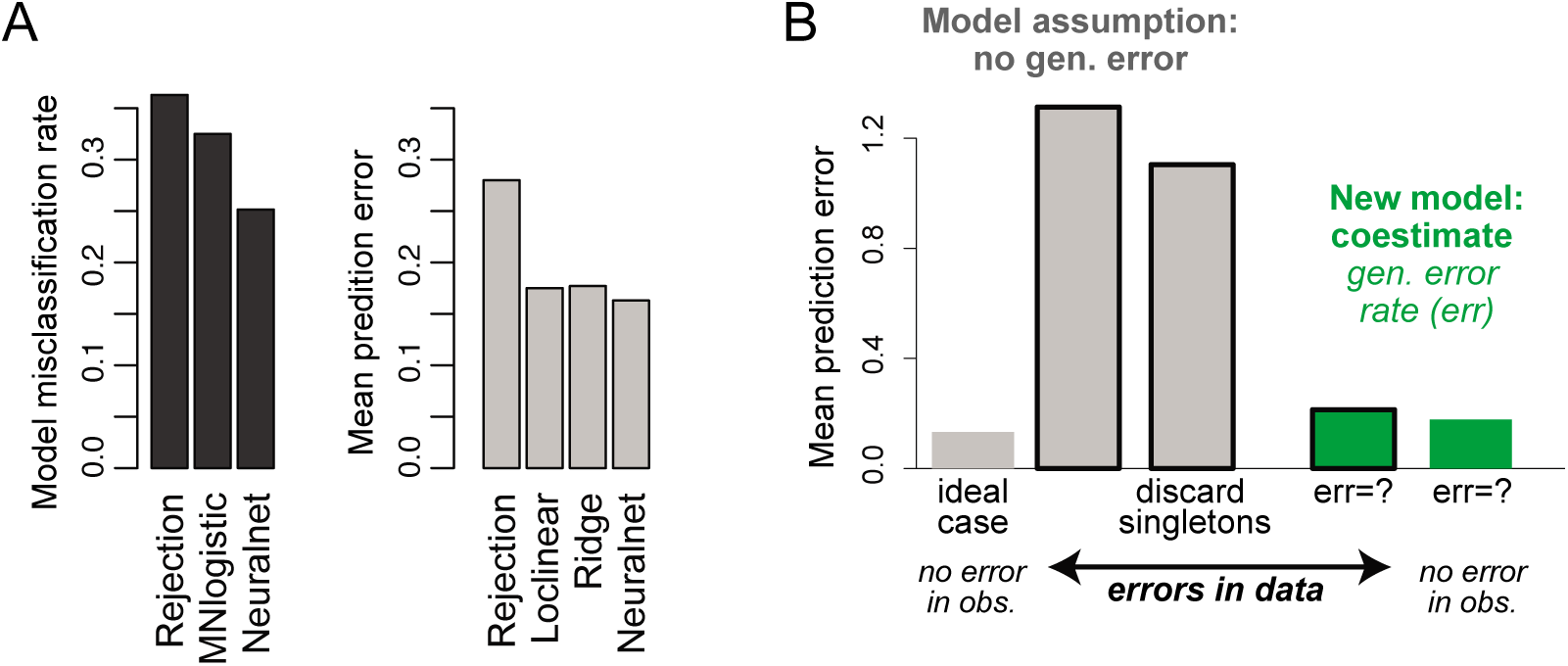
ABC algorithms benchmarked using cross-validation. **A.** ABC routines. Left: Misclassification rates for model selection procedure using rejection, multinomial logistic regression or neural networks. Right: Prediction error for parameter estimation using rejection, local linear regression, ridge logistic regression or neural networks. Errors were averaged over scenarios and parameters. **B.** Impact of genotyping error on previous (ABC neuralnet assuming no error) and new algorithm (ABC neuralnet co-estimating the error rate). Parameter inference was performed on data sets with or without errors, pruned of singletons or not. Prediction errors were averaged over all demographic parameters for the scenario with one expansion.

Given these preliminary results we used algorithms based on neural networks both for parameter estimation and model selection throughout the rest of the study.

In a second phase, we tested the effect of changing the tolerance rate, i.e. the acceptance rate of *pseudo-datasets* for which the statistics were the closest from the observed values, from 0.001 (stringent) to 0.01 and 0.1 (less stringent), for different combination of summary statistics (supplementary fig. 1). Changing the tolerance rate had only a slight impact on the prediction error, with the rate 0.01 usually giving the most accurate estimations. However, it is striking that for some combinations of summaries, the 90% credible intervals were not well estimated for a tolerance of 0.001. The empirical coverages indicated that the interval lengths were underestimated, a non-conservative behavior meaning that they contained the true parameter values less than 90% of the time. The *factor 2* was overall slightly better for tolerance of 0.01 compared to 0.001 while the bias was not clearly impacted by the tolerance rate. For a tolerance of 0.1 the prediction error was not consistently lower than for 0.01, and because the running time was longer, we set the tolerance to 0.01 for all subsequent parameter estimation analyses.

### Genotyping errors

The main advantage of NGS data over SNP data is obviously the fact that a large number of positions are covered along the genome and that we have access to both short and long haplotypic information. Moreover, whole-genome sequencing allows circumventing the well-known issue of SNP ascertainment bias. Nevertheless, the error rate is indisputably higher for NGS than for SNP data and might blur, if not distort, the picture. Ingenious genotype callers (Martin et al. 2010; DePristo et al. 2011; Li 2011; Liao et al. 2017) aim at filling this gap but the difference is still noticeable.

To gauge how these errors impact ABC inference, we generated 1,000 *pseudo-observed datasets* simulated under a simple exponential expansion scenario with errors randomly introduced along chromosomes. We estimated the demographic parameters for each of these pseudo-observed datasets using an ABC procedure based on 300,000 *reference simulations* without any sequencing error. The average prediction error when using all summary statistics was 1.31, which is almost 10 times higher than the average prediction error for 1,000 pseudo-observed datasets without sequencing errors (0.13) (fig. 2B).

A common practice to reduce the impact of sequencing and genotyping errors is to prune the data by ignoring singletons, as they are indeed more likely to be the result of errors than alleles observed at higher frequencies. However, ignoring singletons both in our pseudo-observed datasets and reference simulations only decreased the prediction error to 1.10 (fig. 2B).

We then approximated the imperfect sequencing, genotyping and phasing processes by artificially introducing errors in the reference simulations used in the ABC procedure (see M&M). Using these new reference simulations, we could infer simultaneously the error rate and the demographic parameters. Doing so, we decreased the prediction error rate to 0.21, only 1.6 times higher than the ideal case without errors (fig. 2B). Moreover, when the pseudo-observed data contained no error but the reference simulations did, the average prediction error was relatively close to the ideal case (0.18).

### Five demographic scenarios

We defined five demographic scenarios of interest for human populations but also for other species that might have experienced expansion processes in the past: *One expansion, Two expansions, Bottleneck* (+instant recovery), *Bottleneck and one expansion, Bottleneck and two expansions* (fig. 1). Prior distributions were either uniform or log-uniform distributions (supplementary table S1), with maximal values of 5,000 generations before present for the expansion times and of 500,000 for the population sizes. For the 16 different combinations of summary statistics and two pruning filters (“no pruning” or “singleton pruning”), we evaluated the model misclassification rates and the prediction errors in parameter estimation – as well as several quality criteria such as empirical coverages, estimation biases and factor 2 scores, through intensive cross-validation. For both pruning filters, the genotyping error was co-estimated.

First, we established that the minimum misclassification rate when comparing simultaneously the five models was obtained for the combination of all categories of summary statistics computed once the singletons were removed (supplementary fig. S2, using neural networks and a tolerance rate of 0.001). The misclassification rate was of 0.335, much lower than the 0.8 expected by chance. When trying an easier task, assigning to A: “One or Two expansions” or B: “Bottleneck” or C: “Bottleneck with one or two expansions”, this rate decreased to 0.134. These values partly reflect the fact that some models are nested and that differences between scenarios under two distinct models can be extremely subtle (see next section).

Second, we identified the combination of statistics leading to the best parameter estimation for each of the five scenarios independently. This was done by minimizing the prediction error averaged over all demographic parameters (i.e. all parameters but the genotyping error rate). No combination provided systematically the most accurate estimations, although the AFIBS category was almost always part of the best combination (table 2). Using the best subsets, the prediction error was on average 0.219 (across the five models). Chosen subsets performed well according to other performance criteria, such as empirical coverages, *factor2*, and estimation biases (supplementary fig. S3).

**Table 2.**
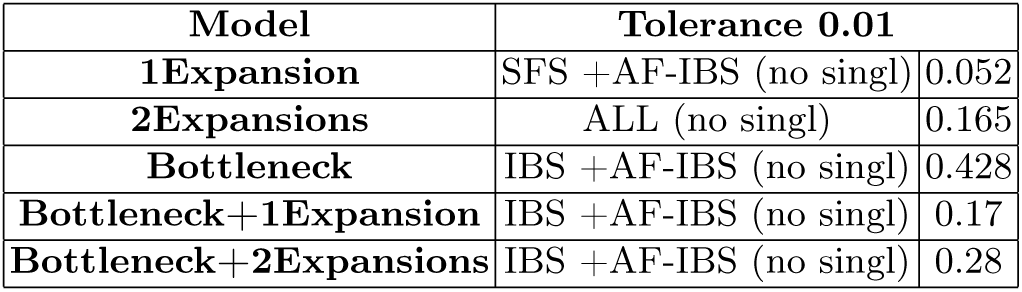
Best combinations of summary statistics for five demographic models. For each model, prediction errors were evaluated on a validation set of 500 pseudo-datasets and averaged over all demographic parameters. Best combinations were identified as the ones with the smallest prediction error. Minimal values are given in the right column. no singl = singletons were pruned.

| Model | Tolerance 0.01 |  |
| --- | --- | --- |
| **1Expansion** | SFS +AF-IBS (no singl) | 0.052 |
| **2Expansions** | ALL (no singl) | 0.165 |
| **Bottleneck** | IBS +AF-IBS (no singl) | 0.428 |
| **Bottleneck+1Expansion** | IBS +AF-IBS (no singl) | 0.17 |
| **Bottleneck+2Expansions** | IBS +AF-IBS (no singl) | 0.28 |

Interestingly the genotyping error rate could be estimated quite accurately along with the demographic parameters. For almost all scenarios and tolerance rates, combining SFS and AFIBS provided the most accurate estimation of this error rate. The only exception was the *Bottleneck* scenario for which AFIBS+IBS performed slightly better when the tolerance rate was 0.01. For this genotyping error rate parameter, the prediction error averaged across the five scenarios was of 0.11 when the tolerance was set to 0.01.

### Detecting two subsequent expansions

When trying to assign a pseudo-observed dataset generated under a two-expansion scenario, to either the one-expansion or the two-expansion model, there is a risk of misclassification inherent to the fact that these models are nested. As depicted on Figure 3 (left), the probability of identifying the model *One expansion* while in fact *Two expansions* occurred increased when the ancient and recent growth rates were very similar and when the old expansion was too weak compared to the recent expansion. More generally two expansions were identified when the ancient growth rate was larger than 1×10^−3^, or between ∼3.5×10^−4^ and 1×10^−3^ if the recent rate was larger (supplementary fig. 4, top left). Under scenarios with bottleneck, the bottleneck strength had an additional impact on misclassification: datasets generated under a *Bottleneck and two expansions* scenario were more commonly assigned to *Bottleneck and one expansion* if the bottleneck was strong and recent (coefficient less than ∼0.2 and time more recent than ∼1000 generations) (fig. 3 middle). Interestingly, even when followed by two expansions, a bottleneck was detectable as long as its coefficient was smaller than ∼0.5 (fig. 3 right).

**Fig. 3.**
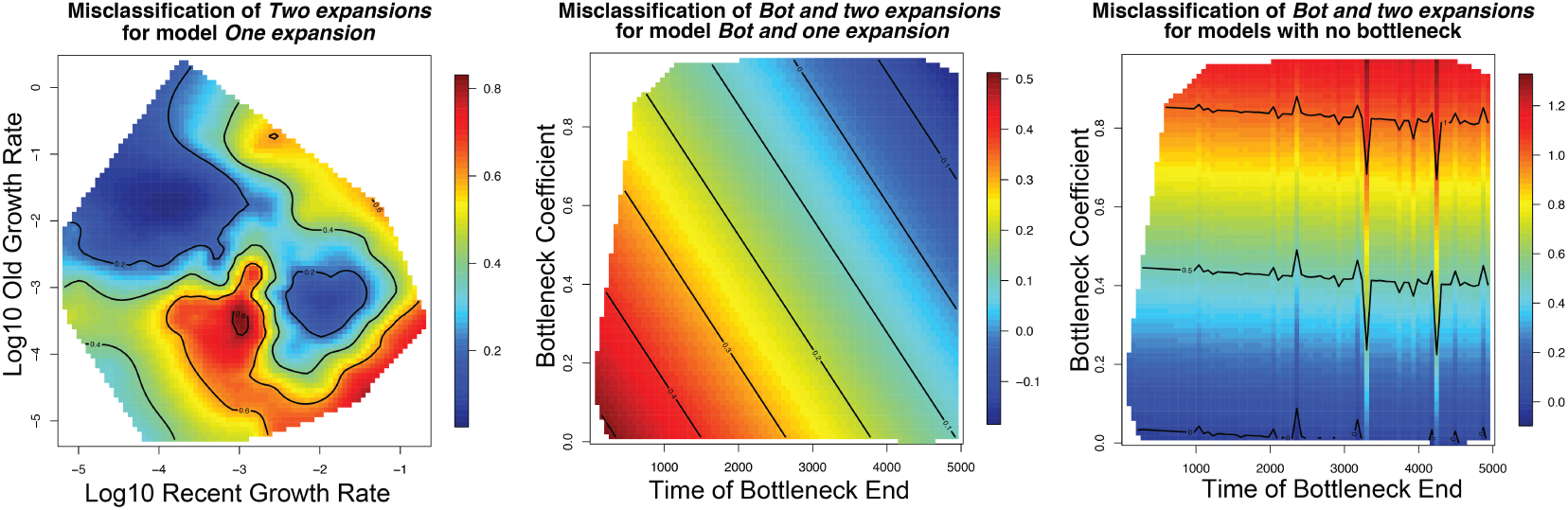
Model selection with nested models. Each of the 2500 validation datasets was assigned to one of the five models using ABC (Fig 1). Misclassification rates were interpolated across different demographic parameters. Left and middle: models with two expansions identified as models with one expansion. Right: models with a bottleneck identified as models without bottleneck. Red denotes highest rates of misclassification and blue lowest rates of misclassification. Interpolations were made using kriging models (R package fields).

**Fig. 4.**
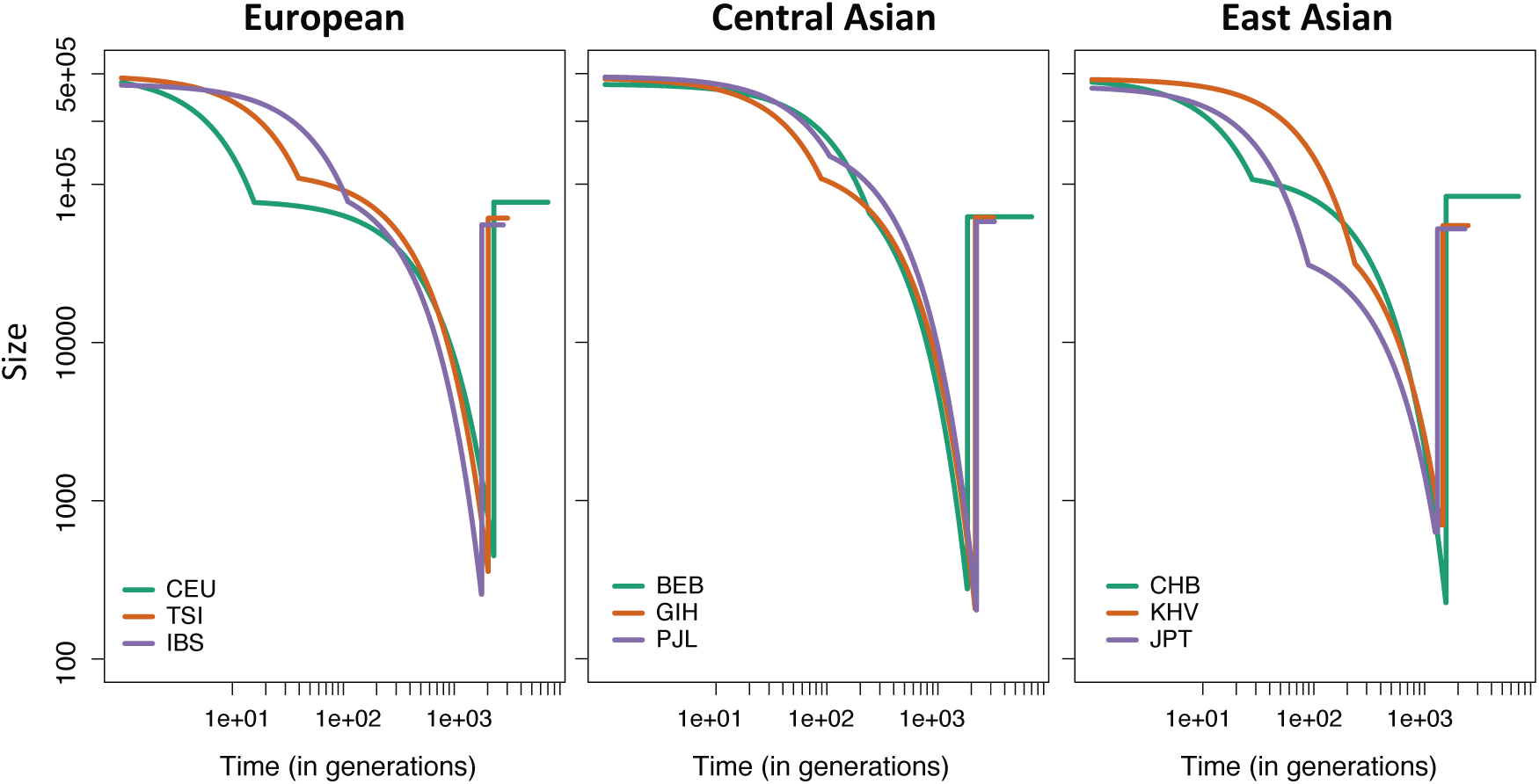
Demographic history inferred with ABC for nine human populations. From Left to right: European (CEU, TSI, IBS), Central Asian (BEB, GIH, PJL), East Asian (CHB, KHV, JPT). For each population, our ABC was run on a randomly subsampled dataset of nine individuals from the 1000 Genomes database. *Bottleneck and two expansions* was independently identified as the best model for each of these populations. The summaries giving the best performance within this model were used (AF-IBS + IBS).

### Application to human data

We applied the ABC analysis to individuals across two datasets: Complete Genomics (CG) and the 1000 Genomes Project (1000g), randomly sampled from multiple groups across Western Africa (Gambian, Mende), Western/Central Africa (Esan, Yoruba), East Africa (Luhya), Europe (Utah residents of European ancestry, Iberian, Tuscan), South Asia (Bengali, Gujarati, Punjabi), and East Asia (Han Chinese, CHB; Kinh, KHV; Japanese, JPT).

### Demography in European, South and East Asian groups

We first performed model selection on all populations and systematically identified the bottleneck followed by two expansions as the most likely scenario for non-African populations. Posterior probabilities associated with this model ranged from 0.78 to 1 (table 3). Overall, the reconstructed demographic histories were quite similar across European, South Asian and East Asian populations sequenced by 1000g. We found evidence for a strong bottleneck, with an estimated reduction in size ranging between 0.0027 and 0.0126 (average 0.006), starting around 1869 generations ago on average across all populations. The bottleneck was followed by an ancient mild expansion with a population size reaching on average 84,205 (average growth rate = 0.003), and by a very recent and strong expansion that led to a current effective population size of 460,392 (ave. growth rate = 0.031). The drastic change in growth rate occurred around 106 generations ago (fig. 4 and table 3). We also inferred the parameters using the smaller tolerance rate of 0.001, which was found to give similar prediction errors but less accurate bounds for the 90% credible intervals (based on cross-validations, see above). In that case, the estimated timing of the recent expansion and the bottleneck were slightly older (166 and 2322 gen. ago respectively, while the estimated current population size was lower (105,239 individuals). We stress out that even if the point estimates for the current population size differ to some extent, a recent strong expansion was found in both cases (recent growth rate of 0.020 versus ancient rate of 0.003).

**Table 3.**
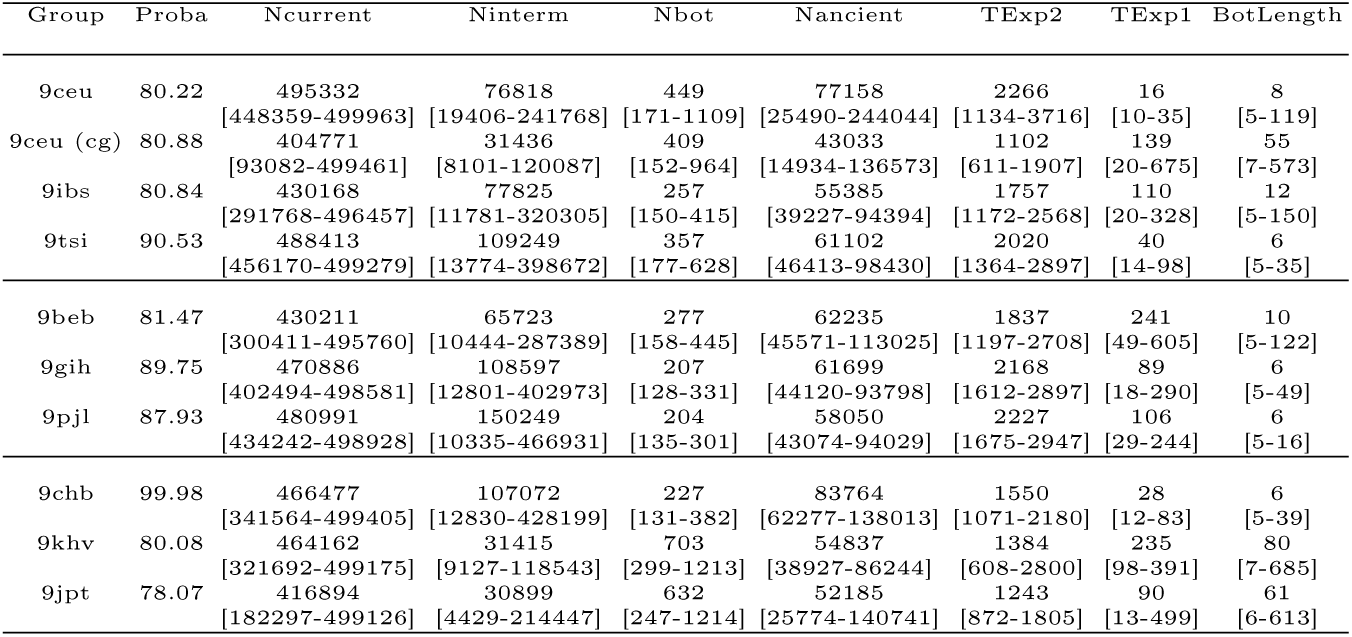
Parameter estimations for Eurasian populations. The column *Proba* displays the posterior probability of the selected model (*Bottleneck and two expansions*). For each model parameter, the median of the posterior distribution is given on the first line and the 95% confidence interval on the second line.

When inferring the scenario with the combination of summary statistics that ranked second in term of prediction error, results were similar (recent expansion 193 gen ago, bottleneck 1575 gen ago, recent growth rates usually larger than the old ones) (supplementary table S3). Recent growth rates were estimated to be higher for East Asians [0.023-0.057], then West Asians [0.012-0.02], and finally Europeans [0.002-0.011]).

We then evaluated the goodness-of-fit of the inferred scenarios using a posterior predictive check approach by checking if the observed summary statistics were in the range of the summary statistics computed from the posterior predictive simulations (ie from data generated under the inferred demographic scenario, see Method). We found that the fit was excellent in most of the European and Asian populations: posterior predictive summary statistics encompassed the observed value for at least 97% of the summary statistics used for parameter inference and at least 76 % the summary statistics not used for parameter inference (eg. SFS, LD, Tajima D, …) (table 4). The fit was slightly poorer in the Iberian, Punjabi and Gujarati populations, where these numbers dropped to 80% for the used summary statistics and 62% for the new ones (averaged over the three populations).

**Table 4.** Goodness-of-fit scores of inferred demographic scenarios evaluated through posterior predictive checks. For each population of the 1000g subset, we report the percentage of observed summary statistics that are in the range of the newly simulated summary statistics, for (a) the statistics already used in the ABC for parameter inference, (b) the remaining summary statistics.

| Group | esn | gwd | lwk | msl | yri | ceu | ibs | tsi | beb | gih | pjl | chb | khv | jpt |
| --- | --- | --- | --- | --- | --- | --- | --- | --- | --- | --- | --- | --- | --- | --- |
| Used Stats (a) | 0.88 | 0.75 | 0.96 | 0.46 | 0.93 | 1.00 | 0.83 | 1.00 | 1.00 | 0.79 | 0.79 | 1.00 | 1.00 | 1.00 |
| New Stats (b) | 0.64 | 0.68 | 0.34 | 0.19 | 0.36 | 0.87 | 0.64 | 0.87 | 0.82 | 0.56 | 0.68 | 0.83 | 0.97 | 0.99 |

### Demographic inference for African groups

When analysing each African population independently, we found that the model selected via ABC for each of these populations was the model assuming a bottleneck and a single expansion (probabilities 0.55-0.99). Moreover, for all five populations, we found that the bottleneck was very recent and strong (409 generations ago on average, effective population size reduced by a factor 0.0095 on average), and was followed by a drastic expansion (average exponential growth rate 0.036) (supplementary table S4). In addition, the bottleneck was systematically estimated to have lasted five generations, which corresponds to the lower bound of the prior distribution. Again, posterior predictive check was performed to further validate the estimated histories. For the Mende, Luhya and Yoruba populations the number of statistics that could be correctly predicted was extremely low: respectively 46%, 96% and 93% of the statistic used for parameter inference, but only 19%, 34% and 36% of the remaining ones (table 4). The fit was slightly better for the Gambian (75% and 68%) and for the Esan (88% and 64%) but still lower than for most Eurasian populations. Among the five African populations, the Gambian and Esan had the most ancient estimates of expansion time (971 and 471 generations ago respectively) and slower growth rates (0.008 and 0.016 resp.).

### Estimating the genotyping error rate

Using the combination of statistics that provided the best estimation of the genotyping error rate (SFS + AFIBS), we estimated that it was larger in the Complete Genomics than in the 1000genomes dataset for both the Utah residents of European ancestry (CEU) and the Yoruba (YRI) *(resp. 8.81 ×10^−6^ and 20 ×10^−6^ in CG versus 5.88 ×10^−6^ and 18.2 ×10^−6^ in 1000G)*. Another striking result was that, while the estimated error rates were reasonably low for all non African populations (below 8.81×10^−6^), they were systematically larger for the African datasets (supplementary fig. S5). While genotyping procedures might differ slightly across populations, such a large discrepancy is unlikely to reflect only a higher rate of errors in the genotype calling for African versus non-African samples. Additionally, this discrepancy cannot rebsult from differences in inference accuracy between demographic models. Indeed, the prediction error of this parameter, as computed using cross-validations, is of the same order of magnitude for both *Bottleneck and one expansion* and *Bottleneck and two expansions* scenarios (0.025 and 0.04). On the other hand, it could reflect the fact that the assumed demographic scenarios fit better the Eurasian than the African data. In the case of poor fit, increasing the error rate offers extra flexibility to explain the data by increasing the noise.

## Discussion

### ABC, neural network, and dimension

The algorithm based on neural networks was the best of all investigated methods. Thanks to its non-linear approach, this method emerges as the best method for handling a large number of summary statistics. This is quite important, since when using a very large number of summary statistics it is fundamental to handle the different kind of information provided by NGS data (such as SFS, LD, length of IBS tracts) to perform valid inferences of both ancient and recent events. Even when taking into account only one category of summary statistics (eg. LD or SFS), we were still considering simultaneously dozens of statistics, and this number reached hundreds when combining all statistics.

ABC suffers from the curse of dimensionality (Blum 2010), so using more and more statistics does not necessarily increase its performance (in particular if these statistics provide redundant information) because we cannot generate an infinite number of pseudo-datasets. The neural network method proposed by Blum and François (2010) acts mainly as a dimension reduction step applied to the high dimensional summary statistics sets. It allows us to increase the tolerance rate and thus to accept many more simulations, alleviating the curse. The large number of accepted simulations is corrected by learning the non-linear relationships that link summary statistics and parameters. In this process, the summary statistics are projected into a space (the hidden layer of the net) of dimension equal to the number of parameters; in this study, this yields a severe reduction in dimension as the parameter space is much smaller than the summary space. Other techniques of dimension reduction have been proposed as a step of ABC (Blum et al. 2013 for review; Pudlo et al. 2016). In a recent paper Sheehan and Song (2016) implemented a deep neural network to bypass the ABC rejection step and detect selective signals in 100kb-regions, which corresponds somehow to setting the tolerance rate to 1 in our method, although the neural network architecture, summary statistics, scenarios of interest and inference algorithms differ. It will be interesting to compare these techniques in future work.

### Sequencing, genotyping, and phasing errors

To reduce the impact of sequencing and genotyping errors we pruned the data by removing singletons but observed that it was not a satisfying solution. This is consistent with findings that “minor-allele-frequency filters – usual practice for genotype-by-sequencing data – negatively affected nearly all estimates” (Shafer et al. 2014). On the other hand, our strategy of modeling genotyping errors via an extra parameter that we estimated together with the demographic parameters provided enough flexibility to estimate correctly the demographic histories of populations, based on samples of variable and unknown quality.

Moreover, we found in our cross-validation study that the error rate could be correctly estimated provided that appropriate summary statistics were used, especially the site-frequency-spectrum along with the allele-frequency IBS statistics (Theunert et al. 2012). It is thus tempting to compare the rates estimated for Complete Genomics (CG) and 1000 genomes (1000g). Yet, the interpretation is not trivial. CG data have a much higher coverage than the 1000g (∼ 51-89x versus ∼ 2-4x), however this might not necessarily translate into a higher quality of CG data for several reasons: (1) The genotypes are called based on likelihoods that take into account other individuals in the dataset. Therefore, the large sample size of the 1000g should counterbalance somehow its low coverage; (2) The same reasoning holds for phasing, which will be more accurate for large datasets, such as 1000g, than for small ones, such as CG and, (3) The sequencing platforms differ between datasets (CG versus Illumina) and are known to have different error rates. For all these reasons, predicting which dataset would have the higher estimated error rate might be dubious. Nevertheless, we found that error rates were generally low and that CG data had higher error rates than 1000g, in concordance with Wall et al. (2014) that estimated the genotype error rates to be higher for sequences obtained by the Complete Genomics technology than by Illumina.

### European and Asian histories. Bottleneck and two successive expansions

For all populations of European and Asian ancestry we detected the Out-Of-Africa bottleneck followed by two successive expansions, an ancient and mild one followed by a recent and stronger one. The ancient expansion was estimated to have started on average 1,190 generations ago (ie 35,700 ya assuming a generation length of 30 years, 95% CI ave. upper bound: 81,728 ya), and the recent expansion 106 generations ago (i.e 3,180 years ago, 95% CI ave. upper bound: 8,460 ya). The most ancient expansion appears thus as a signal of a Paleolithic expansion, while the more recent expansion is consistent with the Neolithic transitions that emerged from 11,500 to 3,500 years ago across the world (Bellwood et al 2005).

This is of interest as only few studies had been able to detect simultaneously both expansions. For example, Aimé et al. (2013; 2014; 2017) used independent datasets (mitochondrial and autosomal short sequences, Y-chromosome and autosomal microsatellite loci) to capture two different time scales and show evidence for expansions before and after the emergence of farming. Palamara et al. (2012) inferred several growth phases in Ashkenazi Jews history using IBD tract lengths, however they focused only on very recent history (less than 200 generations). Following on, Carmi et al. (2014) inferred separately ancient (paleolithic) and recent (medieval) expansions by applying independently a method based on SFS for ancient times and the one based on IBD tract lengths for recent times. Among the few model-based studies that infer simultaneously both expansions, one relies on the SFS-based method *dadi (Gutenkunst et al. 2009)* but uses more than 1000 individuals and all the parameters linked to events preceding the expansions, such as ancient bottlenecks and sizes, were fixed (Tennessen et al. 2012). Similarly, Gazave et al. (2014) used a very large sample of high quality to estimate the very recent growth rate of human populations. Compared to these studies, our approach focuses on datasets with lower sample sizes and quality, and enables reconstructing the broad picture of bottlenecks and growths from a limited amount of data. Remarkably our estimate of the recent growth rate (0.031 averaged across Eurasian populations) is similar to the ones inferred by Tennessen et al. (2012) and Gazave et al. (2014) (0.0307 and 0.0338).

Unlike in African populations, the posterior predictive checks (PPC) performed well for a wide range of summary statistics observed for European and Asian populations. Although there is likely extra complexity in Eurasian demographic histories, the out-of-Africa bottleneck was probably so strong that it wiped out part of the ancient demographic signals in non-African populations compared to African populations. The assumption of a single ancestral population of constant size may thus have a smaller impact on non-African population demographic inference, although we expect some parameter estimates to be biased. For example, the inferred bottleneck could be a compromise between two events identified previously as causing bottlenecks: the Out-Of-Africa migration and the split between European and Asian populations (Keinan et al. 2007; Schaffner et al. 2005). This would lead to an underestimation of the Out-Of-Africa timing.

### Demographic histories in Africa and model--based approach

Our study highlighted that the simple demographic scenarios, still often used to depict African populations, were not able to explain the observed data. Interestingly, it showed that the co-estimated genotyping error rate provides a convenient flag to detect models with poor fit even before performing the posterior predictive check. This rate was indeed consistently larger for datasets that could not be fitted by any proposed model. Note that when we estimated the parameters of the *Bottleneck and one expansion* scenario based on classical statistics (heterozygosity, Tajima’s D, …) and the site-frequency-spectrum, we identified a weak bottleneck (average coefficient 0.75) around 3714 generations ago followed by a mild expansion (average growth rate = 4.4×10^−4^) (supplementary table S5). This history is more similar to what is usually inferred for African populations, such as the Yoruba. Yet, this scenario is not favored when exploiting numerous linkage-informed statistics. We suggest that although this scenario is useful to have a broad picture of past African histories, there is an additional demographic signal that can be picked when extending the summary statistics sets. Part of this complexity was highlighted by various studies based on more comprehensive data (see Schlebusch and Jakobsson 2018 for a recent review). This complexity might encompass additional bottlenecks in the ancient past, complex fluctuations in size or admixture between populations.

Our results advocate for checking thoroughly the goodness-of-fit of inferred models based on posterior checks, and for investigating more complex scenarios that could fit the numerous summary statistics of African data. In particular ancient structure in Africa might be one of the causes, and previous studies (e.g. Sjödin et al. 2012) could be revisited with these new summary statistics in future work.

More generally, the goodness-of-fit of the demographic histories inferred from SMC, IBD or AFS based approaches is too rarely investigated, although the outcome of these approaches is known to be strongly affected by past population structure or complex migration scenarios (see Mazet et al. 2016 for the case of PSMC). Several recent studies illustrated that a demographic scenario proposed based on one specific subset of summary statistics, could be clearly rejected when considering another one (Chikhi et al. 1/2018; Beichman et al. 2017; Lapierre et al. 2017). As demonstrated in this study, ABC is a promising approach in this context, because it provides a natural framework to perform such goodness-of-fit tests.

### Possible methodological extensions

#### Mutation and recombination

In this study we used fixed mutation and recombination rates, set to values commonly assumed in human genetics (see Ségurel et al. 2014 for review) Although it would be hard to co-estimate the mutation rate because mutational effect balances out with the population size (but additional ancient DNA samples could help to disentangle both), the average recombination rate could be co-estimated as done by Boitard et al. (2016). Pseudo-datasets could also be simulated with recombination rates varying along the chromosome, since such maps have been established for human populations with some degree of uncertainty (Kong et al. 2002; Frazer et al. 2007; Hinch et al. 2011). The first solution should be beneficial mostly for non-model species for which even a rough average rate is unknown, while the second solution could be tested for humans. However, adjusting the recombination maps according to the population of interest and simulating data tailored for each genomic region would lead to a substantial increase in the number of simulations required.

#### Neutrality

When studying human demography, given the current knowledge of the genetic map composition, it is possible to discard presumed non-neutral regions such as protein-coding regions, human accelerated regions (HAR), DNA hypersensitive sites (DHS), and others (see Schraiber and Akey 2015 for review). However doing so would prevent us from extracting continuous 2-MB long regions from the real data. As it is assumed in most demographic inference methods (PSMC, MSMC, IBS-based approaches, …), given that such non-neutral regions are uncommon, we considered here that their impact on the overall demographic signal would be limited. However, several studies have demonstrated the strong confounding effects of background selection. Future developments are needed to address the challenging task of co-estimating selection and demography (Bank et al., 2014; Ewing and Jensen, 2016; Hernandez et al., 2011; McVicker et al., 2009; Sheehan and Song, 2016).

## Conclusion

We implemented an ABC method that infers population demography from sequencing data, while accounting for the specificities of these data (correlation between close SNPs, genotyping errors …). We demonstrated that this method allows inferring demographic histories consisting of successive bottleneck and expansions from a limited number of individuals. We successfully applied this method to human populations from Eurasia, inferring the out-of-Africa bottleneck and two successive expansions corresponding to the Paleolithic and Neolithic periods.

An additional goal of our study was to understand what commonly used population genetic summary statistics should an *ABC for sequencing data* framework rely on. We conclude that it depends on the investigated demographic model, but that the allele-frequency identity-by-state (Theunert et al, 2012) is a key statistic that consistently increased prediction accuracy.

Even though more and more sequences are available, it is likely that dataset will remain of limited size in the coming years for most non-model species. Therefore, methods that make the best of a sample of intermediate size are much needed, and our approach will be useful for studying populations in many species.

## Material and methods

### Simulated data

We simulated sequences of length 2Mbp using *fastsimcoal 2.1 (Excoffier et al. 2013).* This program generates quickly long neutral sequences with recombination based on the Sequential Markov coalescent *(McVean and Cardin 2005; Marjoram and Wall 2006)*. For each parameter set, we simulated 100 independent tracts, with a constant mutation rate of 1.25×10^−8^ per site per generation, and a constant recombination rate of 1×10^−8^ per site per generation (Scally and Durbin 2012; Schiffels and Durbin 2014). We generated around 300,000 pseudo-datasets for each of the five demographic models displayed on Figure 1. They consist in one optional bottleneck followed either by an instant recovery, or by one or two expansions. These exponential expansions happen in the last 5,000 generations, and the population sizes can range from 100 to 500,000 (see supplementary table S1 for detailed information on parameters and priors).

For preliminary analyses, simulations were generated under models of constant size followed by either one or two exponential expansions with narrower priors (the sizes were constrained to be under 70,000).

When taking sequencing errors into account, we introduced an extra parameter, the *error rate*, associated with a uniform prior distribution with bounds [0, 2×10^−5^]. At each position in the initial sample (even monomorphic positions) and for each haplotype, the allele was switched from derived to ancestral – or from ancestral to derived – with a probability equal to this error rate.

### Summary statistics

For each dataset of 100 independent 2Mb-long regions sequenced for *n* diploid individuals we computed the following statistics, grouped into five categories.

#### Classic

1. Proportion of segregating sites over the genome (total number of SNPs *S* divided by the total sequencing length = 200 Mb)

2. Tajima’s D statistic averaged over regions

3. Expected heterozygosity at a segregating site computed

4. Average haplotypic heterozygosity along the genome. Haplotypic heterozygosities were computed for non-overlapping windows of size 50-kb as 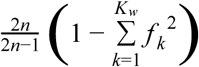, where *K*_*w*_ is the number of unique haplotypes in a given window *w*, and *f*_*k*_ is the frequency of the *k*-th haplotype.

*SFS*

5. Unfolded site-frequency-spectrum (percentage of segregating sites for which the derived allele frequency is *i*, for all i in *[1, …, 2n-1])*, and the total proportion of segregating sites (already used in “Classic”). 2n summary statistics.

6. Statistics linked to the variability of these counts at each bin [1,…, 2n-1]. These were computed for each allele frequency bin, as the standard deviation of the distances separating two adjacent SNPs at frequency *i*. 2n-1 summary statistics.

*LD*

7. The linkage disequilibrium is computed as the average *r*^*2*^ for pairs of SNPs. Values were stratified by the physical distance separating SNPs. Following Boitard et al (2016), we considered 19 bins of distances (for which the mean ranged from 282 bp to 1.4 Mb). 19 summary statistics.

*IBS*

8. The length distributions of identical-by-state segments shared between *m* haplotypes were summarized by 10 percentiles. Those were computed for m=2, 4, 8 and 16 haplotypes. IBS segments were defined as regions of the genome completely identical across haplotypes. 40 summary statistics.

*AF-IBS*

9. At each SNP, the AF-IBS segment is identified as the region extending around this SNP and identical across all haplotypes carrying the derived allele at the focal SNP (Theunert et al. 2012). The lengths of AF-IBS segments are stratified by the frequency at their focal SNP, leading to a distribution for each frequency [2/n,… (2n-1)/n]. We summarize each distribution by its mean and variance. 2*(2n-1) summary statistics.

All summary statistics were computed using our custom python scripts that will be available online.

### Approximate Bayesian Computation (ABC)

ABC is a Bayesian framework designed for models with unknown likelihood but under which one can generate data through computer simulations. It aims at selecting the model that best explains the observed data among several possible scenarios, as well at estimating the parameters of this model. Given a distribution *a priori* of the parameters, ABC approximates their posterior distribution via two simplifying steps: (1) the full dataset is reduced to a set of *summary statistics*, (2) the new posterior given these *summary statistics* is obtained through the inspection of numerous pseudo-datasets. Those pseudo-datasets are simulated using the generative model with parameter values drawn from specified prior distributions. Pseudo-datasets for which summary statistics are close enough to the observed ones are *accepted* and their corresponding parameter values are considered as a sample of the approximate posterior distribution (standard *rejection* algorithm). In refined ABC algorithms these *accepted* pseudo-datasets are used to learn a local relation linking the summary statistics to the model parameters, this assumed relation being: a linear model (Beaumont et al. 2002), a linear model penalizing large coefficients to better account for collinearity between summary statistics (ridge regression, Blum et al. 2013), or a non-linear model (calibrated using feed-forward neural networks, Blum and François 2010). Formally, given *S* the observed summary statistics, θ the true parameter, *S*_*i*_ the summary statistics of pseudo-dataset *i* generated with parameter values θ_*i*_, and η the tolerance error, a model f is learned from accepted simulations so that for all i, θ _*i*_ = *f* (*S*_*i*_) + ε_*i*_ with ε _*i*_ a random variable with mean zero and constant variance (in homoscedastic models). Accepted parameter values are then adjusted as followed, 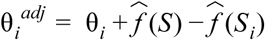 and weighted by the Euclidean distance between *S* and *S*_*i*_, to approximate the posterior distribution. For more details and formal descriptions of heteroscedastic models (where the variance is not constant) see Blum and François (2010).

As recommended by Blum and François (2010), before learning the different models, we applied a logit transformation to the parameters with bounds corresponding to the bounds of their prior distribution.

When running the leave-one-out cross-validations, the prediction error was calculated as

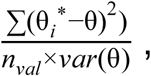

where θ is the true parameter value, θ* is the predicted parameter value, and nval is the number of points where true and predicted values are compared (Csilléry et al. 06/2012).

We also computed the relative bias as

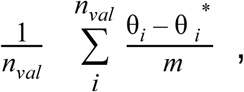

where, 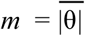, the average of the absolute values of θ. and the *factor 2* as the proportion of test datasets for which the point estimate is at least half and at most twice the true value, i.e.

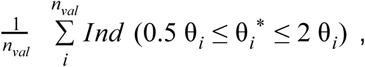

where *Ind* denotes the indicator function (Cornuet et al. 2008).

All ABC analyses were performed with the R package ‘abc’ (Csilléry et al. 2012) and we tested four methods for parameter estimation: rejection, linear regression, ridge regression, neural network, and three methods for model selection that compute posterior model probabilities (rejection, multinomial logistic regression, neural network).

### Publicly available genomes

#### Complete Genomics

We used a dataset of 54 unrelated individuals present in the panel “69 Genomes” from Complete Genomics (Drmanac et al. 2010). The data phased by Rasmussen et al (2014) using SHAPEIT2 (Delaneau et al. 2013) are available at compgen.cshl.edu/ARGweaver/CG_results/download/. Our ABC procedure was applied independently to nine individuals of European ancestry (Utah residents, CEU) and 9 individuals from the African Yoruba population (YRI). Applied filters are described in the supplementary material.

#### 1000 genomes

The latest version of the 1000g phase 3 data was downloaded from ftp.1000genomes.ebi.ac.uk (release 20130502, file date 20140730). These genomes were already phased using an improved version of SHAPEIT2 (Delaneau et al. 2014; Consortium 2015) Once again we applied our ABC approach to nine individuals chosen randomly from each of the following groups: African (Esan, ESN; Gambian, GWD; Luhya, LWK; Mende, MSL; Yoruba, YRI), European (Utah residents of European ancestry, CEU; Iberian, IBS, Tuscan, TSI), South Asian (Bengali, BEB; Gujarati, GIH; Punjabi, PJL), and East Asian (Han Chinese, CHB; Kinh, KHV; Japanese, JPT). Applied filters are described in the supplementary material.

## Acknowledgments

This work was supported by the ANR grant DemoChips (12-BSV7-0012) coordinated by FA, which funded in particular the salary of FJ, and by the NEFREX project funded by the European Union (People Marie Curie Actions, International Research Staff Exchange Scheme, call FP7-PEOPLE-2012-IRSES-number 318979). Large scale simulations were ran on the Genotoul bioinformatics platform Toulouse Midi-Pyrénées (www.bioinfo.genotoul.fr). We thank Bertrand Servin for sharing his original python code, Michael Blum for discussions about the ‘abc’ R package, and Trevor Pemberton for discussions about error rates in CG and 1000G datasets.

## References

Aimé C, Austerlitz F. 2017. Different kinds of genetic markers permit inference of Paleolithic and Neolithic expansions in humans. Eur. J. Hum. Genet. 25:360–365.

Aime C, Laval G, Patin E, Verdu P, Segurel L, Chaix R, Hegay T, Quintana-Murci L, Heyer E, Austerlitz F. 2013. Human Genetic Data Reveal Contrasting Demographic Patterns between Sedentary and Nomadic Populations That Predate the Emergence of Farming. Mol. Biol. Evol. 30:2629–2644.

Aimé C, Verdu P, Ségurel L, Martinez--Cruz B, Hegay T, Heyer E, Austerlitz F. 2014. Microsatellite data show recent demographic expansions in sedentary but not in nomadic human populations in Africa and Eurasia. Eur. J. Hum. Genet. 22:1201–1207.

Batini C, Lopes J, Behar DM, Calafell F, Jorde LB, van der Veen L, Quintana-Murci L, Spedini G, Destro-Bisol G, Comas D. 2011. Insights into the Demographic History of African Pygmies from Complete Mitochondrial Genomes. Mol. Biol. Evol. 28:1099–1110.

Beaumont MA. 2008. Joint determination of topology, divergence time, and immigration in population trees. In: Matsumura S, Forster P, Renfrew C, editors. Simulation, Genetics, and Human Prehistory. Cambridge: McDonald Institute for Archaeological Research. p. 135–154.

Beaumont MA, Zhang W, Balding DJ. 2002. Approximate Bayesian Computation in Population. Genetics 162:2025–2035.

Beichman AC, Phung TN, Lohmueller KE. 2017. Comparison of Single Genome and Allele Frequency Data Reveals Discordant Demographic Histories. G3 7:3605–3620.

Bhaskar A, Song YS. 2014. Descartes’ rule of signs and the identifiability of population demographic models from genomic variation data. Ann. Stat. 42:2469–2493.

Bhaskar A, Wang YXR, Song YS. 2015. Efficient inference of population size histories and locus-specific mutation rates from large-sample genomic variation data. Genome Res. 25:268–279.

Blum MGB. 2010. Approximate Bayesian computation: a nonparametric perspective. J. Am. Stat. Assoc. 105:1178–1187.

Blum MGB, François O. 2010. Non-linear regression models for Approximate Bayesian Computation. Stat. Comput. 20:63–73.

Blum MGB, Nunes MA, Prangle D, Sisson SA. 2013. A Comparative Review of Dimension Reduction Methods in Approximate Bayesian Computation. Stat. Sci. 28:189–208.

Boitard S, Rodríguez W, Jay F, Mona S, Austerlitz F. 2016. Inferring Population Size History from Large Samples of Genome-Wide Molecular Data - An Approximate Bayesian Computation Approach. Beaumont MA, editor. PLOS Genet. 12:e1005877.

Browning SR, Browning BL. 09/2015. Accurate Non-parametric Estimation of Recent Effective Population Size from Segments of Identity by Descent. Am. J. Hum. Genet. 97:404–418.

Carmi S, Hui KY, Kochav E, Liu X, Xue J, Grady F, Guha S, Upadhyay K, Ben-Avraham D, Mukherjee S, et al. 2014. Sequencing an Ashkenazi reference panel supports population-targeted personal genomics and illuminates Jewish and European origins. Nat. Commun. 5:4835.

Chikhi L, Rodríguez W, Grusea S, Santos P, Boitard S, Mazet O. 2018. The IICR (inverse instantaneous coalescence rate) as a summary of genomic diversity: insights into demographic inference and model choice. Heredity 120:13–24.

Consortium T 1000 GP. 2015. A global reference for human genetic variation. Nature 526:68–74.

Cornuet J-M, Santos F, Beaumont MA, Robert CP, Marin J-M, Balding DJ, Guillemaud T, Estoup A. 2008. Inferring population history with DIY ABC: a user-friendly approach to approximate Bayesian computation. Bioinformatics 24:2713.

Cox MP, Morales DA, Woerner AE, Sozanski J, Wall JD, Hammer MF. 2009. Autosomal Resequence Data Reveal Late Stone Age Signals of Population Expansion in Sub-Saharan African Foraging and Farming Populations. PLOS One 4:e6366.

Csilléry K, Blum MGB, Gaggiotti OE, François O. 2010. Approximate Bayesian computation (ABC) in practice. Trends Ecol. Evol. 25:410–418.

Csilléry K, François O, Blum MGB. 06/2012. abc: an R package for approximate Bayesian computation (ABC): R package: abc. Methods Ecol. Evol. 3:475–479.

Delaneau O, Marchini J, Consortium T 1000 GP. 2014. Integrating sequence and array data to create an improved 1000 Genomes Project haplotype reference panel. Nat. Commun. 5, 3934.

Delaneau O, Zagury J-F, Marchini J. 2013. Improved whole-chromosome phasing for disease and population genetic studies. Nat. Methods 10:5–6.

DePristo MA, Banks E, Poplin R, Garimella KV, Maguire JR, Hartl C, Philippakis AA, Angel G del, Rivas MA, Hanna M, et al. 2011. A framework for variation discovery and genotyping using next-generation DNA sequencing data. Nat. Genet. 43:491–498.

Drmanac R, Sparks AB, Callow MJ, Halpern AL, Burns NL, Kermani BG, Carnevali P, Nazarenko I, Nilsen GB, Yeung G, et al. 2010. Human Genome Sequencing Using Unchained Base Reads on Self-Assembling DNA Nanoarrays. Science 327:78–81.

Excoffier L, Dupanloup I, Huerta-Sanchez E, Sousa VC, Foll M. 2013. Robust Demographic Inference from Genomic and SNP Data. Akey JM, editor. PLOS Genet. 9:e1003905.

Excoffier L, Estoup A, Cornuet J-M. 2005. Bayesian Analysis of an Admixture Model With Mutations and Arbitrarily Linked Markers. Genetics 169:1727–1738.

Excoffier L, Schneider S. 1999. Why hunter-gatherer populations do not show signs of Pleistocene demographic expansions. Proc. Natl. Acad. Sci. U. S. A. 96:10597–10602.

Fontaine MC, Snirc A, Frantzis A, Koutrakis E, Öztürk B, Öztürk AA, Austerlitz F. 2012. History of expansion and anthropogenic collapse in a top marine predator of the Black Sea estimated from genetic data. Proc. Natl. Acad. Sci. U. S. A. 109:E2569–E2576.

Frazer KA, Ballinger DG, Cox DR, Hinds DA, Stuve LL, Gibbs RA, Belmont JW, Boudreau A, Hardenbol P, Leal SM, et al. 2007. A second generation human haplotype map of over 3.1 million SNPs. Nature 449:851–861.

Gattepaille LM, Jakobsson M, Blum MGB. 2013. Inferring population size changes with sequence and SNP data: lessons from human bottlenecks. Heredity 110:409–419.

Gazave E, Ma L, Chang D, Coventry A, Gao F, Muzny D, Boerwinkle E, Gibbs RA, Sing CF, Clark AG, et al. 2014. Neutral genomic regions refine models of recent rapid human population growth. Proceedings of the National Academy of Sciences 111:757–762.

Gutenkunst RN, Hernandez RD, Williamson SH, Bustamante CD. 2009. Inferring the Joint Demographic History of Multiple Populations from Multidimensional SNP Frequency Data. McVean G, editor. PLOS Genet. 5:e1000695.

Harris K, Nielsen R. 2013. Inferring demographic history from a spectrum of shared haplotype lengths. PLOS Genet. 9:e1003521.

Hinch AG, Tandon A, Patterson N, Song Y, Rohland N, Palmer CD, Chen GK, Wang K, Buxbaum SG, Akylbekova EL, et al. 2011. The landscape of recombination in African Americans. Nature 476:170–175.

Keinan A, Mullikin JC, Patterson N, Reich D. 10/2007. Measurement of the human allele frequency spectrum demonstrates greater genetic drift in East Asians than in Europeans. Nat. Genet. 39:1251–1255.

Kong A, Gudbjartsson DF, Sainz J, Jonsdottir GM, Gudjonsson SA, Richardsson B, Sigurdardottir S, Barnard J, Hallbeck B, Masson G, et al. 2002. A high-resolution recombination map of the human genome. Nat. Genet. 31:241–247.

Lapierre M, Lambert A, Achaz G. 2017. Accuracy of demographic inferences from the site frequency spectrum: the case of the Yoruba population. Genetics 206:439–449.

Liao P, Satten GA, Hu Y-J. 2017. PhredEM: a phred-score-informed genotype-calling approach for next-generation sequencing studies. Genet. Epidemiol. 41:375–387.

Li H. 2011. A statistical framework for SNP calling, mutation discovery, association mapping and population genetical parameter estimation from sequencing data. Bioinformatics 27:2987–2993.

Li H, Durbin R. 2011. Inference of human population history from individual whole-genome sequences. Nature 475:493–496.

Li S, Jakobsson M. 2012. Estimating demographic parameters from large-scale population genomic data using Approximate Bayesian Computation. BMC Genet. 13:22.

Liu X, Fu Y-X. 2015. Exploring population size changes using SNP frequency spectra. Nat. Genet. 47:555–559.

MacLeod IM, Larkin DM, Lewin HA, Hayes BJ, Goddard ME. 2013. Inferring Demography from Runs of Homozygosity in Whole-Genome Sequence, with Correction for Sequence Errors. Mol. Biol. Evol. 30:2209–2223.

Marjoram P, Wall JD. 2006. Fast” coalescent” simulation. BMC Genet. 7:1.

Martin ER, Kinnamon DD, Schmidt MA, Powell EH, Zuchner S, Morris RW. 2010. SeqEM: an adaptive genotype-calling approach for next-generation sequencing studies. Bioinformatics 26:2803–2810.

Mazet O, Rodriguez W, Grusea S, Boitard S, Chikhi L. 2016. On the importance of being structured: instantaneous coalescence rates and human evolution—lessons for ancestral population size inference? Heredity 116:362–371.

McVean GAT, Cardin NJ. 2005. Approximating the coalescent with recombination. Philos. Trans. R. Soc. Lond. B Biol. Sci. 360:1387–1393.

Palamara PF, Lencz T, Darvasi A, Pe’er I. 2012. Length Distributions of Identity by Descent Reveal Fine-Scale Demographic History. Am. J. Hum. Genet. 91:809–822.

Palstra FP, Heyer E, Austerlitz F. 2015. Statistical Inference on Genetic Data Reveals the Complex Demographic History of Human Populations in Central Asia. Mol. Biol. Evol. 32:1411–1424.

Patin E, Laval G, Barreiro LB, Salas A, Semino O, Santachiara-Benerecetti S, Kidd KK, Kidd JR, Veen LV der, Hombert J-M, et al. 2009. Inferring the Demographic History of African Farmers and Pygmy Hunter–Gatherers Using a Multilocus Resequencing Data Set. PLOS Genet. 5:e1000448.

Patin E, Siddle KJ, Laval G, Quach H, Harmant C, Becker N, Froment A, Régnault B, Lemée L, Gravel S, et al. 2014. The impact of agricultural emergence on the genetic history of African rainforest hunter-gatherers and agriculturalists. Nat. Commun. 5, 3163.

Pudlo P, Marin J-M, Estoup A, Cornuet J-M, Gautier M, Robert CP. 2016. Reliable ABC model choice via random forests. Bioinformatics 32:859–866.

Rasmussen MD, Hubisz MJ, Gronau I, Siepel A. 2014. Genome-wide inference of ancestral recombination graphs. PLOS Genet. 10:e1004342.

Scally A, Durbin R. 2012. Revising the human mutation rate: implications for understanding human evolution. Nat. Rev. Genet. 13:745–753.

Schaffner SF, Foo C, Gabriel S, Reich D, Daly MJ, Altshuler D. 2005. Calibrating a coalescent simulation of human genome sequence variation. Genome Res. 15:1576.

Schiffels S, Durbin R. 2014. Inferring human population size and separation history from multiple genome sequences. Nat. Genet. 46:919–925.

Schlebusch CM, Jakobsson M. 2018. Tales of Human Migration, Admixture, and Selection in Africa. Annu. Rev. Genomics Hum. Genet. 19:405–428.

Schraiber JG, Akey JM. 2015. Methods and models for unravelling human evolutionary history. Nat. Rev. Genet. 16:727.

Ségurel L, Wyman MJ, Przeworski M. 2014. Determinants of Mutation Rate Variation in the Human Germline. Annu. Rev. Genomics Hum. Genet. 15:47–70.

Shafer ABA, Gattepaille LM, Stewart REA, Wolf JBW. 2014. Demographic inferences using short-read genomic data in an Approximate Bayesian Computation framework: in silico evaluation of power, biases, and proof of concept in Atlantic walrus. Mol. Ecol. 24: 328–345.

Sheehan S, Harris K, Song YS. 2013. Estimating variable effective population sizes from multiple genomes: a sequentially markov conditional sampling distribution approach. Genetics 194:647–662.

Sheehan S, Song YS. 2016. Deep learning for population genetic inference. PLOS Comput. Biol. 12:e1004845.

Sjödin P, Agnès E. Sjöstrand, Jakobsson M, Blum MGB. 2012. Resequencing Data Provide No Evidence for a Human Bottleneck in Africa during the Penultimate Glacial Period. Mol. Biol. Evol. 29:1851–1860.

Soares P, Alshamali F, Pereira JB, Fernandes V, Silva NM, Afonso C, Costa MD, Musilová E, Macaulay V, Richards MB, et al. 2012. The Expansion of mtDNA Haplogroup L3 within and out of Africa. Mol. Biol. Evol. 29:915–927.

Sunnåker M, Busetto AG, Numminen E, Corander J, Foll M, Dessimoz C. 2013. Approximate Bayesian Computation. Wodak S, editor. PLOS Comput. Biol. 9:e1002803.

Tennessen JA, Bigham AW, O’Connor TD, Fu W, Kenny EE, Gravel S, McGee S, Do R, Liu X, Jun G, et al. 2012. Evolution and Functional Impact of Rare Coding Variation from Deep Sequencing of Human Exomes. Science 337:64–69.

Terhorst J, Kamm JA, Song YS. 2017. Robust and scalable inference of population history from hundreds of unphased whole genomes. Nat. Genet. 49:303–309.

Theunert C, Tang K, Lachmann M, Hu S, Stoneking M. 2012. Inferring the History of Population Size Change from Genome-Wide SNP Data. Mol. Biol. Evol. 29:3653–3667.

Veeramah KR, Hammer MF. 2014. The impact of whole-genome sequencing on the reconstruction of human population history. Nat. Rev. Genet. 15:149–162.

Wall JD, Tang LF, Zerbe B, Kvale MN, Kwok P-Y, Schaefer C, Risch N. 11/2014. Estimating genotype error rates from high-coverage next-generation sequence data. Genome Res. 24:1734–1739.

Wollstein A, Lao O, Becker C, Brauer S, Trent RJ, Nürnberg P, Stoneking M, Kayser M. 2010. Demographic History of Oceania Inferred from Genome-wide Data. Curr. Biol. 20:1983–1992.

